# Animacy and real world size shape object representations in the human medial temporal lobes

**DOI:** 10.1101/304824

**Authors:** Anna Blumenthal, Bobby Stojanoski, Chris Martin, Rhodri Cusack, Stefan Köhler

**Affiliations:** department of Psychology, University of Western Ontario, London, Ontario; The Brain and Mind Institute, University of Western Ontario, London, Ontario; Department of Psychology, University of Toronto, Toronto, Ontario; Department of Psychology, Trinity College, Dublin, Ireland; Rotman Research Institute, Baycrest, Toronto, Canada

## Abstract

Identifying what an object is, and whether an object has been encountered before, is a crucial aspect of human behavior. Despite this importance, we do not have a complete understanding of the neural basis of these abilities. Investigations into the neural organization of human object representations have revealed category specific organization in the ventral visual stream in perceptual tasks. Interestingly, these categories fall within broader domains of organization, with distinctions between animate, inanimate large, and inanimate small objects. While there is some evidence for category specific effects in the medial temporal lobe (MTL), it is currently unclear whether domain level organization is also present across these structures. To this end, we used fMRI with a continuous recognition memory task. Stimuli were images of objects from several different categories, which were either animate or inanimate, or large or small within the inanimate domain. We employed representational similarity analysis (RSA) to test the hypothesis that object-evoked responses in MTL structures during recognition-memory judgments also show evidence for domain-level organization along both dimensions. Our data support this hypothesis. Specifically, object representations were shaped by either animacy, real-world size, or both, in perirhinal and parahippocampal cortex, as well as the hippocampus. While sensitivity to these dimensions differed when structures when probed individually, hinting at interesting links to functional differentiation, similarities in organization across MTL structures were more prominent overall. These results argue for continuity in the organization of object representations in the ventral visual stream and the MTL.

## INTRODUCTION

The ability to identify the objects we encounter in our daily lives, and know which ones we have seen before, is a crucial aspect of human behavior. What type of object a ‘thing’ is, and whether it is familiar or novel, can drastically change how we might interact with it, including, for example, whether to approach or avoid it. Despite the importance of object recognition, and the relative fluidity with which most humans perform it, a detailed understanding of the neural functional architecture that supports this ability is still elusive. One promising approach to understanding the neural architecture of object perception and memory is to explore how object representations are organized. In particular, it is possible to examine similarities between patterns of brain activity that different types of objects evoke, and to map this neural representational geometry to relevant dimensions in perception and behavior.

It is known that some correspondence exists between how objects are represented in the brain and how we behaviorally categorize them. Important insight has been gained from functional magnetic resonance imaging (fMRI) investigations of object processing in the ventral visual stream (VVS). Numerous fMRI studies have revealed regions within the VVS that preferentially respond to particular stimulus categories with high ecological relevance, including faces, scenes, bodies, and words (see Op de Beeck et al., 2008, for review). Specifically, in extrastriate cortex, regions have been reported that prefer one of these categories over other categories, such as the fusiform face area or the parahippocampal place area (Kanwisher et al., 1997; Epstein & Kanwisher, 1998; Downing et al., 2001; for a review see Kanwisher & Dilks, 2013). Interestingly, these functionally circumscribed regions are systematically organized within broader preference zones. Medial aspects of occipito-temporal cortex typically show a preference for inanimate objects, whereas lateral aspects show a preference for animate objects (Martin, 2007; Grill-Spector & Weiner, 2014; Sha et al., 2015). In addition to the animacy dimension, a number of fMRI studies have revealed large-scale organization of the VVS by real-world size (Konkle & Caramazza, 2013; Konkle, et al., 2012; Cusack et al., 2016). It has been found that there is a preference zone for large inanimate objects in medial occipito-temporal cortex and for small inanimate objects in more dorsolateral aspects, but no corresponding size-based distinction has been found for animate objects in lateral occipito-temporal cortex. This pattern of preferences has been referred to as a tripartite organizing schema (Konkle & Oliva, 2012; Konkle & Caramazza, 2013).

Evidence for organization by animacy and real world size has also come from studies based on multivariate pattern analyses of fMRI data. In this analytical approach, activity is not averaged across voxels, but the similarity between patterns of activity evoked by different stimuli or within a given region are compared. If stimuli within a category evoke more similar patterns of activity than stimuli from different categories, the brain region is considered to contain representations of that category. Inasmuch as the pattern of activity across voxels can be labeled a neural representation of an object, one can think of the comparisons between categories as now existing in “representational geometry”. Interestingly, Kriegeskorte et al. (2008) applied this approach across using voxels distributed throughout the ventral temporal cortex for a wide variety of objects, and found a highly consistent category– and domain-based organization, with evidence for a distinction between animate and inanimate objects, as well as varying degrees of similarity between categories within these domains (see also Proklova, Kaiser, & Peelen, 2016). A recent fMRI study with a similar focus on representational similarities has shown that real-world size is also an organizing dimension of objects across a large swath of temporo-parieto-occipital cortex, as well as within a number of subregions across the VVS (Julian et al., 2016).

The influence of the category and domain of objects has been most thoroughly characterized in the posterior and lateral aspects of the VVS. At present, evidence that speaks to the organization of object representations in medial temporal lobe (MTL) structures is more limited. While it has long been known that memory functions pertaining to objects rely on the integrity of MTL structures (Graham et al., 2010; Eichenbaum et al. 2007; Davachi, 2006), the organization of object representation that supports these functions remains incompletely understood at present. Furthermore, little is known about similarities and differences in organization across different MTL structures, including perirhinal cortex (PrC), parahippocampal cortex (PhC), and the hippocampus (HiP). The more posterior aspect of the PhC has been well characterized, given that it comprises a significant proportion of the parahippocampal place area, a functionally-defined region that preferentially responds to scenes and large objects with navigational relevance (Aguirre et al., 1998; Epstein and Kanwisher, 1998; Epstein & Vass, 2014; Troiani et al., 2012; Konkle & Oliva, 2012). However, it is less clear whether this characterization holds for PhC as a whole, and, whether it also holds for PrC and HiP. The lack of evidence is surprising, given that a more anterior structure in the parahippocampal gyrus, namely, PrC has been proposed to be the apex of the VVS (Murray and Bussey, 1999; Bussey et al., 2007). Furthermore, evidence from studies of structural connectivity in non-human primates as well as functional connectivity studies in humans indicate that both PhC and PrC have strong connectivity with downstream areas in the VVS and other posterior cortical regions (see Ranganath & Ritchey, 2012, for a review). As such, it remains an unknown but interesting possibility that the major dimensions that have been shown to shape representations in the posterior VVS, i.e., animacy and real-world size, also shape organization of object categories in PrC and the PhC. To the extent that the HiP receives much of its cortical input from these structures, it also important to include the HiP in this inquiry.

Research with direct comparisons of visual stimulus responses in PrC and PhC has shown robust differences for processing of faces, objects, and scenes across both structures. At the univariate level, PhC shows a scene preference, while PrC, in particular anterior portions, shows a face preference (Liang et al., 2012; Litman et al., 2010; O’Neil et al., 2013; see Collins and Olson, 2014, for review). However, evidence suggests this is not a sharp distinction, but an anterior-posterior gradient from scenes to faces (Liang et al., 2012; Litman et al., 2010). In MVPA based studies, it has been shown that object, scene, and face information can be distinguished at the category level in both PhC and PrC. In general scene decoding is higher in PhC, and face responses can be better decoded from PrC (LaRocque et al., 2013; Huffman & Stark, 2014, Liang et al., 2012), although Diana et al. (2008), did not find above chance decoding of objects or faces in PrC. Aside from the evidence that scenes and faces are distinctly represented in these MTL structures, it is less clear whether other object categories are distinctly represented, and how this is similar or different across regions. This is in large part due to the fact that most studies have used mixed groups of objects without any systematic attempt to probe category based distinctions. In recent work from our lab, Martin et al. (2013; 2016) explored this issue in the context of recognition memory judgments, using chairs, faces, and buildings as categorized stimuli. We reported that it was possible to decode the perceived familiarity of faces from activity patterns in PrC, the familiarity of buildings from patterns in PhC, and familiarity for chairs from patterns in both structures. While these findings go beyond showing a distinctions between scenes and faces in the medial temporal lobe (MTL), they do not allow for a broader characterization of representational space across a wider variety of object categories.

Our primary interest in the present study was in a comparison of object representations across the PrC, PhC, and in the HiP. While previous work has revealed some evidence for object category specificity in PrC and PhC, the HiP has been seen as more “agnostic”, or insensitive to visual stimulus category (Huffman & Stark, 2014; LaRocque et al., 2013; Diana et al., 2009). It has been posited that this is because the HiP binds object and spatial information received from the PrC and PhC (Eichenbaum et al., 2012; Ranganath & Ritchey, 2012). More specifically, if the HiP represents complex conjunctions of many different kinds of objects and the spatial backdrop of those objects, it may be difficult to reveal any category specificity (e.g., in a complex scene there may be objects from many different categories). Interestingly, one study reported above-chance decoding of scene information from posterior HiP (Liang et al., 2012). At the univariate level, the HiP often shows more activity for scenes as compared to other stimulus categories such as objects or faces, which has led to the suggestion that it be considered a part of the core scene-network (Hodgetts et al., 2016). Furthermore, individuals with hippocampal damage have been reported to show impairments in vividly recalling scenes, maintaining scenes in working memory, and constructing scenes in their imagination (Hassabis et al., 2007; Mullally et al., 2012; Addis et al., 2007). This suggests that the HiP may not be entirely agnostic to the nature of stimulus categories encountered. As such, it is possible that it may also be sensitive to stimulus domain.

In the present fMRI study, we addressed whether and how animacy and real-world size affect the organization of object categories in the MTL. We tested the hypothesis that object-evoked responses in perirhinal and parahippocampal cortex, as well as the hippocampus, show evidence for domain-level organization along both dimensions. To this end, we scanned participants while they performed a continuous recognition memory task on objects from 12 different categories. We chose a continuous recognition memory task because it required participants to make memory decisions (i.e., “old” or “new”) for specific exemplars from these categories, thus maximizing the need to disambiguate objects with substantial feature overlap. To address our questions of interest, we employed representational similarity analyses (RSA). With these analyses we first asked whether PrC, PhC, and the HiP represent distinct categories of objects. We then explored whether the categories were organized along an animate/inanimate divide, and whether or not inanimate objects were organized by their real-world size.

## MATERIALS AND METHODS

### Participants

Fifteen individuals participated in the study (20-32 years of age, mean age = 27.5 years; 8 females). All participants were right-handed with normal or corrected-to-normal vision, and no history of psychiatric or neurological disorders. Data from two participants were excluded due to technical difficulties. Participants received financial compensation for their participation, and provided informed consent according to procedures approved by the University of Western Ontario Health Sciences Research Ethics Board

### Stimuli

Stimuli were color images depicting exemplars from 12 different object categories, including 4 categories of animate objects (faces, bodies, monkeys, and insects), 4 categories of large inanimate objects (buildings, vehicles, trees, and furniture), and 4 categories of small inanimate objects (flowers, fruits, musical instruments, and tools). Size and animacy classification was based on prior research (Konkle et al., 2013) and confirmed through ratings in pilot work in a separate group of participants for all stimuli employed here. Twenty-eight objects were chosen from each category, for a total of 336 experimental stimuli. In addition, 3 filler items were presented in each run, one of which was repeated early on in the run to ensure that participants would immediately be prepared for repetitions. The second and third filler items were presented towards the end of the run to increase the proportion of novel stimuli at that stage. Filler items were chosen from categories other than (and unrelated to) those employed on experimental trials. Images of objects were obtained from the Konkle lab database (http://konklab.fas.harvard.edu/#) and through an additional Google image search. Each image was presented in isolation on a white background bound at 500 × 500 pixels, on a uniform grey background. The size of each image was bound at a maximum of 500 × 500 pixels, for one dimension, with the other dimension the corresponding to the appropriate aspect-ratio. Across categories, there were no significant differences in the area covered by objects in the images, their aspect-ratio, or their mean luminance (all p > 0.05).

### Experimental procedure

During fMRI scanning, participants performed a continuous recognition memory task that required recognition of repeated presentation of specific exemplars (Fig. 2). Exemplars were presented twice, with repetitions always occurring in the same run. Images were presented for 1200 ms, and participants were asked to indicate whether the image was “novel” (1st presentation), or “old” (2nd presentation) with button presses using their middle or index finger. To encourage rapid responding and mark the time window for responding, a red border surrounding the image appeared 600 ms after stimulus onset and stayed on screen until stimulus offset. Participants were instructed to respond as soon as the red border appeared. Mapping of responses to buttons was counterbalanced across participants. Each stimulus presentation was followed by a jittered ITI (2000-6000 ms) during which participants viewed a fixation cross centered on a grey background. Jitter was distributed such that the average delay between first and second presentation of items was matched across categories (average time = 84.1 s, range = 19.0-316.0 s). In addition, the average number of images between repetitions was matched across categories (average number of intervening images =17, range =16-18/ Each run consisted of 4 objects from each of the 12 categories, resulting in a total of 8 image presentations per category, or 96 experimental trials per run. In addition, each run contained 3 filler trials. Across runs, presentations of objects from each category were preceded and followed by an object from each of the other categories with roughly equal frequency (8-11 times/ Participants completed seven runs. Three different run orders were created for the purpose of counterbalancing across participants. Prior to scanning, each participant completed a 5-minute practice task with images from categories that were unrelated to those used during scanning in order to be familiarized with task requirements and response deadline.

**Fig 1.**
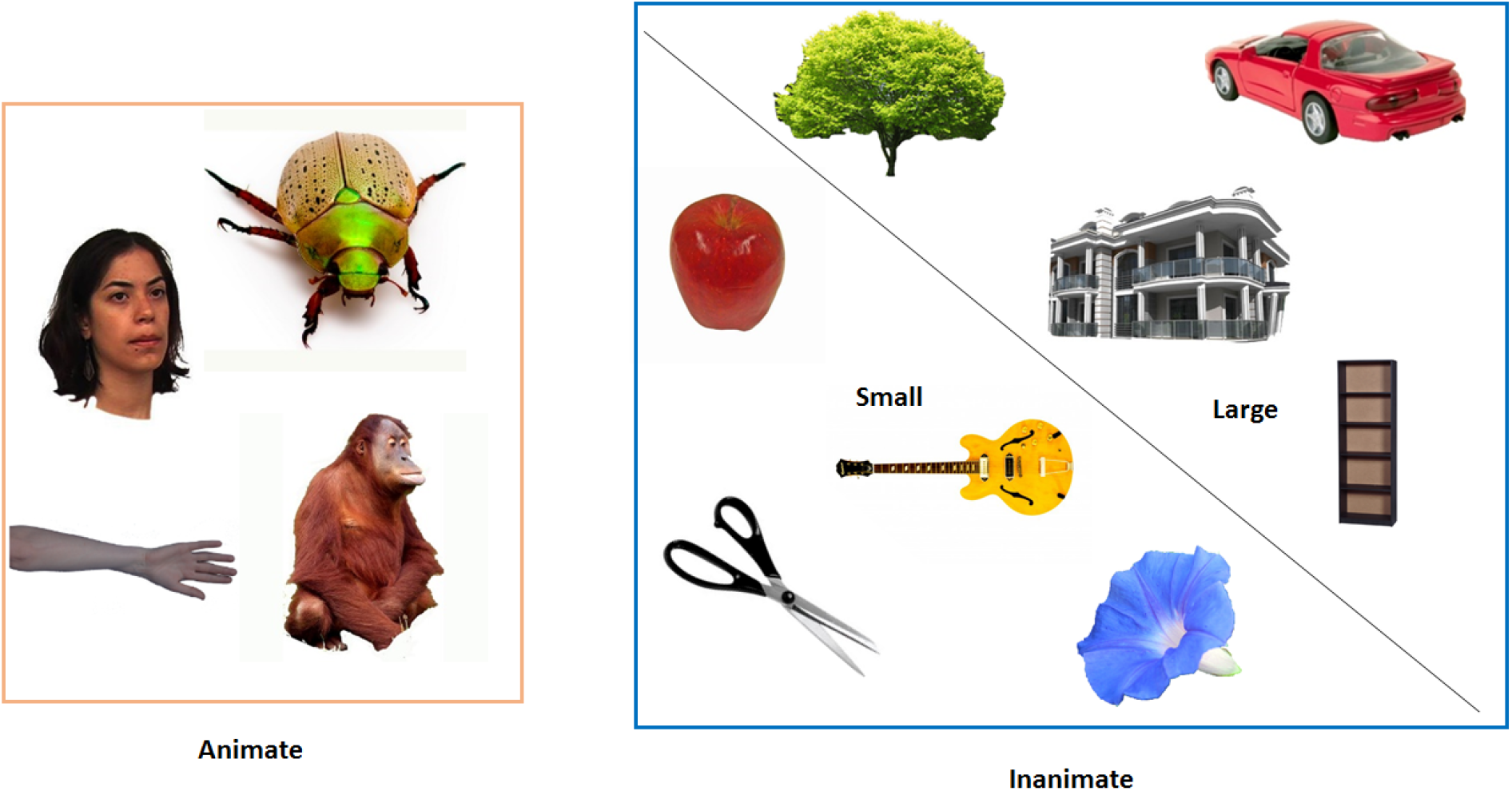
Stimuli. Example objects from the 12 object categories employed. Categories were grouped into animate: faces, bodies, monkeys, insects, inanimate small: flowers, fruits, tools, musical instruments, and inanimate large: buildings, trees, vehicles, furniture.

**Fig 2.**
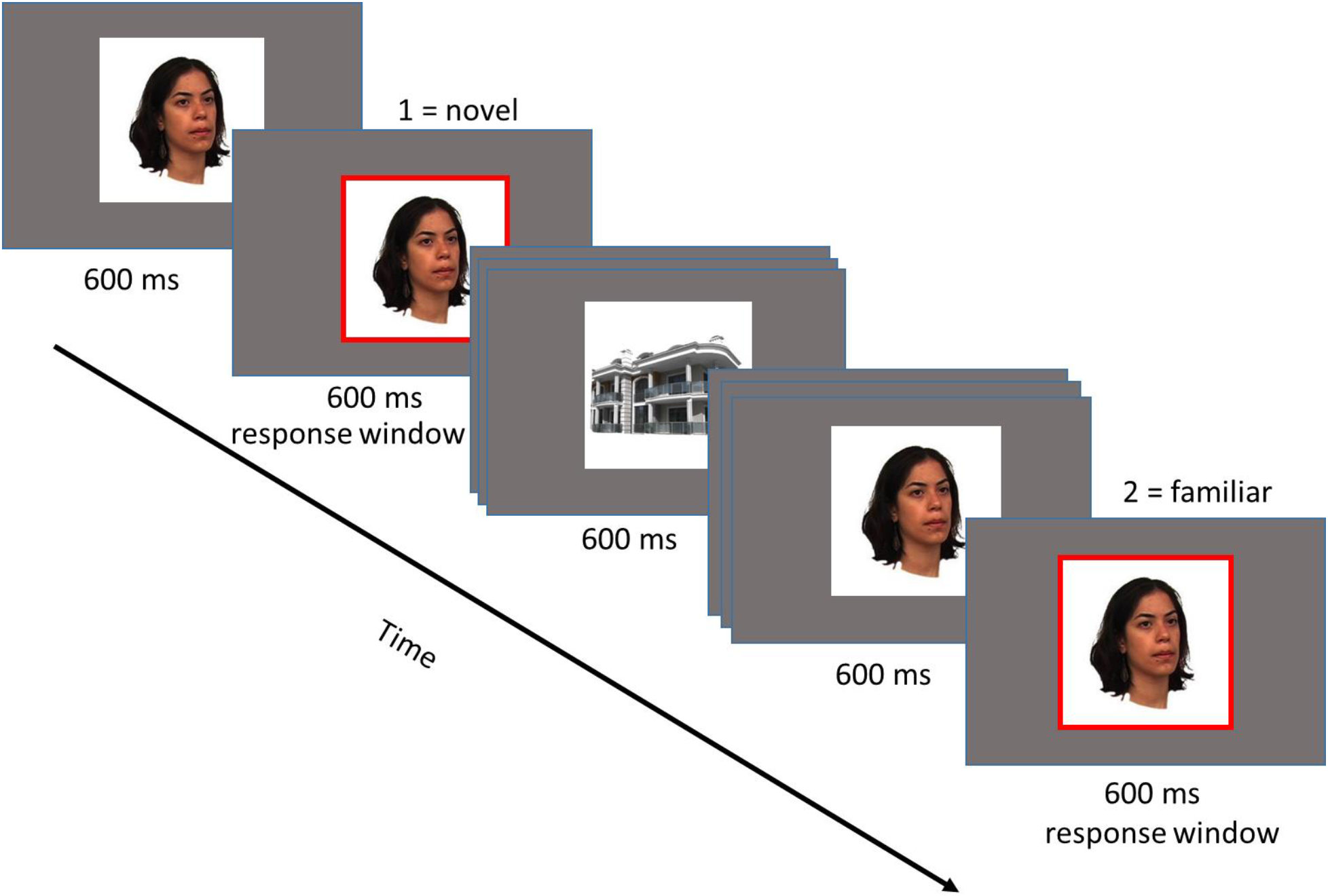
Task: continuous recognition memory. An image depicting an object from one of the twelve categories was presented on screen for 1200ms. After 600ms a red border popped up around the image, and participants were required to respond “novel” indicating it was the first time they had seen that image, or “familiar” indicating that it was the second time they had seen that image.

### Image acquisition

MRI data were acquired on a Siemens TIM Trio 3-Tesla scanner with a high-resolution protocol. Functional MRI volumes were collected using a highly accelerated gradient-echo EPI sequence (Center for Magnetic Resonance Research, University of Minnesota) with a multiband acceleration factor 4 and GRAPPA in-plane acceleration of 2. The following parameters were used: TR=650 ms, TE=30 ms, slice thickness = 2 mm, FOV = 192 mm × 192 mm, flip angle = 54 degree. Each functional volume included 40 slices collected in an interleaved manner. To optimize MR signal in the anterior temporal lobes, a transverse orientation was chosen for acquisition, which allowed for inclusion of the entire temporal and occipital lobes, with partial coverage of frontal and parietal cortices, in all participants. T1-weighted anatomical images were obtained using an ADNI MPRAGE sequence (192 slices, TR = 2300 ms, TE =2.98 ms, 1 mm isotropic voxels, FOV = 240 × 256 mm, flip angle = 9 degrees).

### Neuroimaging analysis

### Preprocessing and Modelling

fMRI data were analyzed using SPM8 (Welcome Institute of Cognitive Neurology; http://www.fil.ion.ucl.ac.uk/spm/software/spm8/), employing an analysis pipeline as implemented in the automatic analysis system (aa)

(www.github.com/rhodricusack/automaticanalysis), (Cusack et al., 2015). Functional data were motion corrected and high pass-filtered to remove low frequency noise (drift); slice-time correction was not implemented due to the use of a multiband sequence. Four dummy scans at the start of each session were discarded to allow for T1 relaxation. For each participant, the mean functional image was then co-registered with the participant-specific anatomical image. Coregistered images were kept in native space for each participant, and no spatial smoothing was applied in order to preserve high-spatial resolution for MVPA. Functional data were convolved using a canonical hemodynamic response function. Categories were modeled, regardless of whether a trial was a 1^st^ or 2^nd^ presentation (12 regressors per run) using a general linear model. Regressors were constructed from boxcars with a duration of each stimulus presentation (1200ms), and were convolved with SPM’s canonical hemodynamic response function. Beta estimates for each category were derived based on 4 exemplars and their repetition in each run. Regressors of no interest included 6 motion regressors. Beta estimates derived from these models were used as input for the univariate and multivariate analyses. Medial temporal lobe ROIs were demarcated manually for each participant on the high-resolution structural images in native space, using the anatomical protocols published by Pruessner et al. (2000; 2002) with adjustments to the posterior border of PhC as specified by Franko et al. (2014). (Fig. 3).

**Fig 3.**
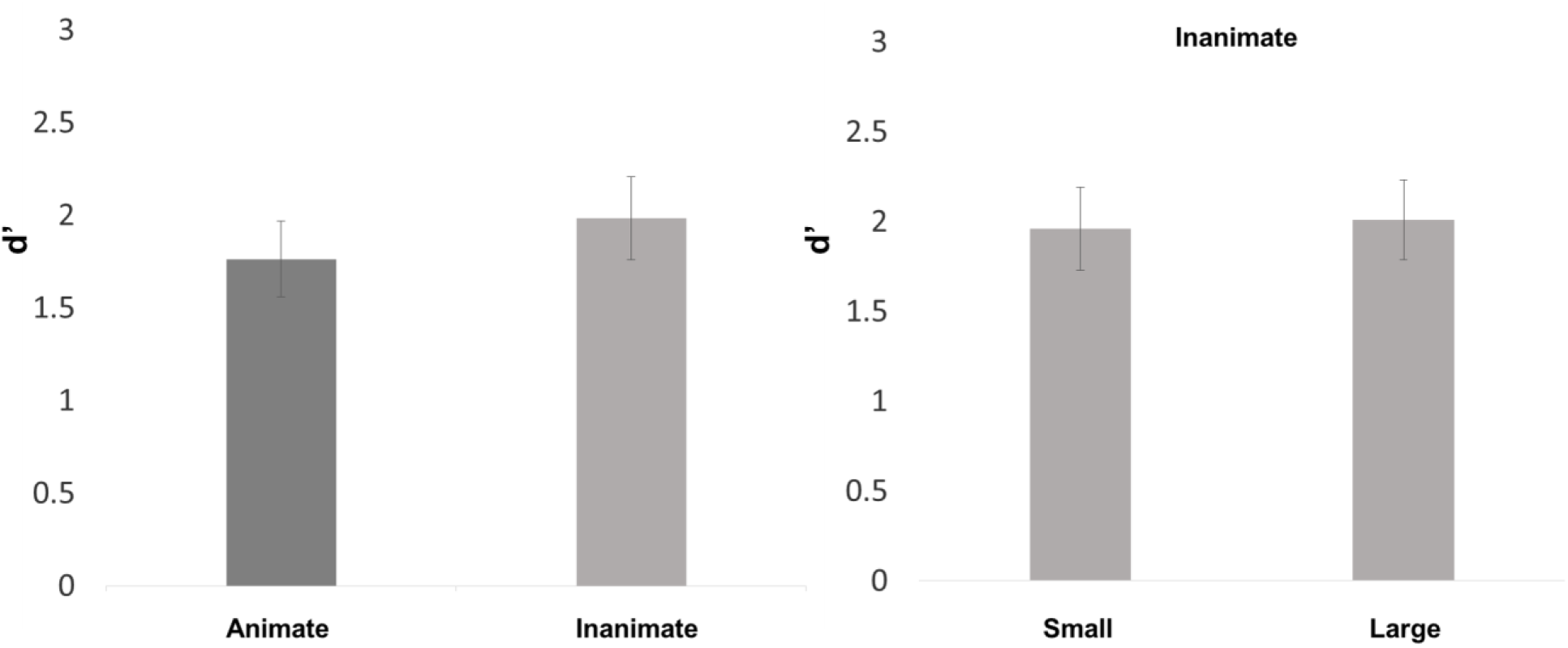
Recognition memory performance for domains of interest, as measured with dprime. There were no significant differences on performance between animate or inanimate objects (p =. 2), or between large and small inanimate objects (p =. 7).

### Univariate analyses

Univariate analyses were conducted for the purpose of selecting features (i.e., voxels) to be included in the multivariate analyses. Towards this end, we contrasted all experimental trials against baseline (gray screen with a fixation cross), which resulted in robust activation throughout occipital and temporal cortex (including MTL) in each participant. We then selected the 20% of voxels with the highest beta values in this contrast (i.e., stimuli > baseline) in each participant-specific anatomically defined ROI. These voxels were used for all remaining multivariate analyses (see Kriegeskorte et al., 2008, 2008b for rationale).

### Representational similarity analysis

Multivariate analyses were computed on a between-run basis to ensure the different comparisons did not vary in temporal proximity (Linke et al., 2011). To explore the representational space in each ROI, for each subject, we first extracted beta values for each category and computed the Pearson’s correlation for each category compared to each other category. Prior to computing the correlations, the grand mean (i.e., the cocktail mean) for each run was subtracted across all voxels for that run (Walther et al. 2015). This resulted in a 12 × 12 representational similarity matrix (RSM) for each participant, for each ROI, with within category similarity values (across runs) on the diagonal, and between category information (across runs) on the off diagonal (Fig 5). To test whether the representational space was modulated by category, animacy, and size within inanimate objects, we created linear models (predefined contrasts) specifying which RSM correlation values were to be subjected to a t-test that tested models (see Fig. 6). These analyses were performed on data in single-subject RSMs, with the group statistics calculated from the average results. For the purpose of visualizing our results, RSMs were then averaged across participants, resulting in a final group similarity matrix for each ROI (Kriegeskorte, Mur, Bandettini, 2008). Group-averaged RSMs were ordered in the following way: animate objects, small inanimate objects, and large inanimate objects. Note that RSM’s are not symmetrical in the visualization, this is because the upper half of the matrix shows the mean from a subset of across run correlations (i.e., cell 1, 2 is condition 1 in the even runs correlated with condition 2 in the odd runs, whereas cell 2, 1, is condition 1 in the odd runs correlated with condition 2 in the even runs).

We first asked whether there was evidence of category-level organization in each ROI. To test for this, we defined a contrast of category representation (see Fig. 6), in other words a linear model where all within category (diagonal) patterns were more highly correlated than between category (off diagonal) patterns. In the initial analysis, we tested an omnibus contrast (i.e., model) that probed for the presence of any category-specific information in each ROI. We then tested for information relating to each of the 12 categories individually. Specifically, we tested whether the patterns of activation across voxels were more similar within each category compared to the 11 other categories, using subject as a random effect.

In our second set of analyses, we asked whether or not the animate vs inanimate object distinction that has been found to shape the organization of object representations in more posterior aspects of the VVS (Konkle & Caramazza, 2013; Konkle, et al., 2012) was also an organizing dimension in the MTL. This analysis was identical to the previously described analyses, except that for the purpose of evaluating differences in correlations (i.e., within vs between) we focused on the domains of animate as compared to inanimate objects rather than individual categories (see Fig. 6). Importantly, in these analyses we removed the diagonal from our model in order to discard the influence of within category similarities.

In our third and final set of analyses, we asked whether real world size is an organizing dimension within the domain of inanimate objects in MTL, again as has been reported for object representations in more posterior aspects of the VVS (Kriegeskorte et al., 2008; Proklova, Kaiser, and Peelen, 2016). Here, we divided inanimate objects into groups of small or large objects, large objects included trees, furniture, vehicles and buildings, and small objects including fruit, flowers, musical instruments, and tools. The analysis was identical to the previous one except that within versus between similarities were computed across all categories of large or small inanimate objects (see Fig. 6). As in the analyses on animacy described above, we did not include the diagonal in testing of this model.

## RESULTS

### Behavioral Results

Recognition-memory accuracy, indexed using the discriminability index *ď*, and reaction times are shown in Table 1 for all categories. Critically, memory discrimination as measured with *d’* was matched across dimensions of interest. Specifically, we found no differences in performance between animate and inanimate objects (Mean *d’* animate = 1.76, *SD*=0.78, Mean *d’* inanimate =1.94, *SD*=0.71, *t(12)* =-1.30, *p* =. 2 (Fig. 4)). There were also no differences based on real-world size, i.e., between large inanimate and small inanimate objects (Mean *d’* large inanimate = 1.96, *SD*=0.77, Mean d’ small inanimate = 2.00, SD=0.80, *t* (12) = −.452,*p* =. 7 (Fig. 4). We did find differences in RTs between animate and inanimate categories (Mean RT animate = 1.01 s, *SD* = 0.037, Mean RT inanimate = 1.00 s *SD* = 0.041 *t*(12) = 2.41, *p* = 0.02), as well as large inanimate and small inanimate objects (Mean RT large inanimate = 1.007 s, *SD* = 0.040, Mean RT small inanimate = 0.993 s, *SD* = 0.033, *t*(12) = −3.49, *p* = 0.004). Although these RT differences are statistically significant, we note that they are very small because the task required responding within a restricted time window (i.e., there was a response deadline that was visually indicated in the displays). We think it is unlikely that differences of this magnitude are reflected in the BOLD response we report, in particular given the focus on patterns of activity that have been de-meaned.

**Fig 4.**
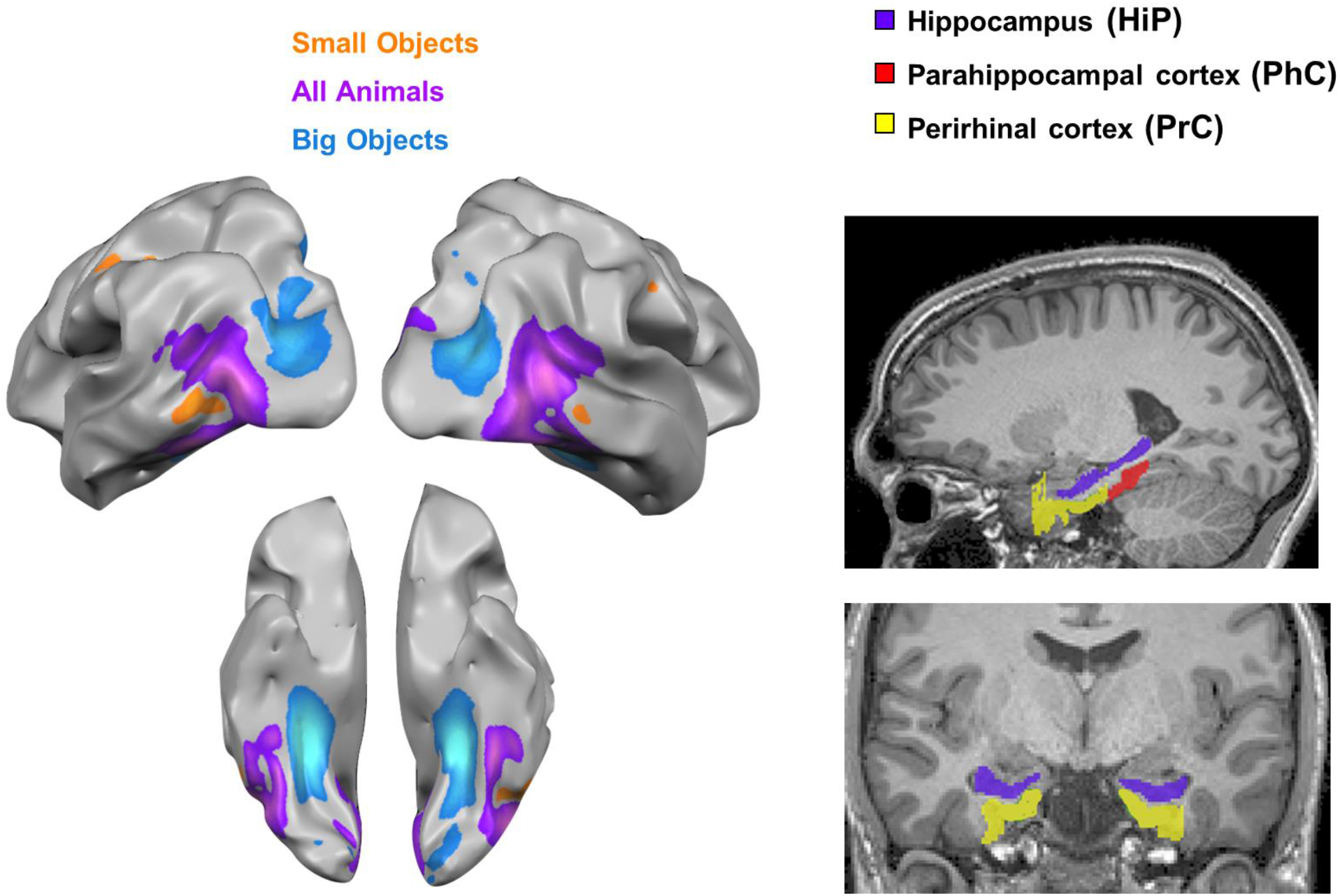
Left: Tripartite preference zones in posterior occipito-temporal cortex (courtesy of Konkle et al.,2013). Right: Anatomical regions of interest examined in the medial temporal lobe, for one example participant.

**Fig 5.**
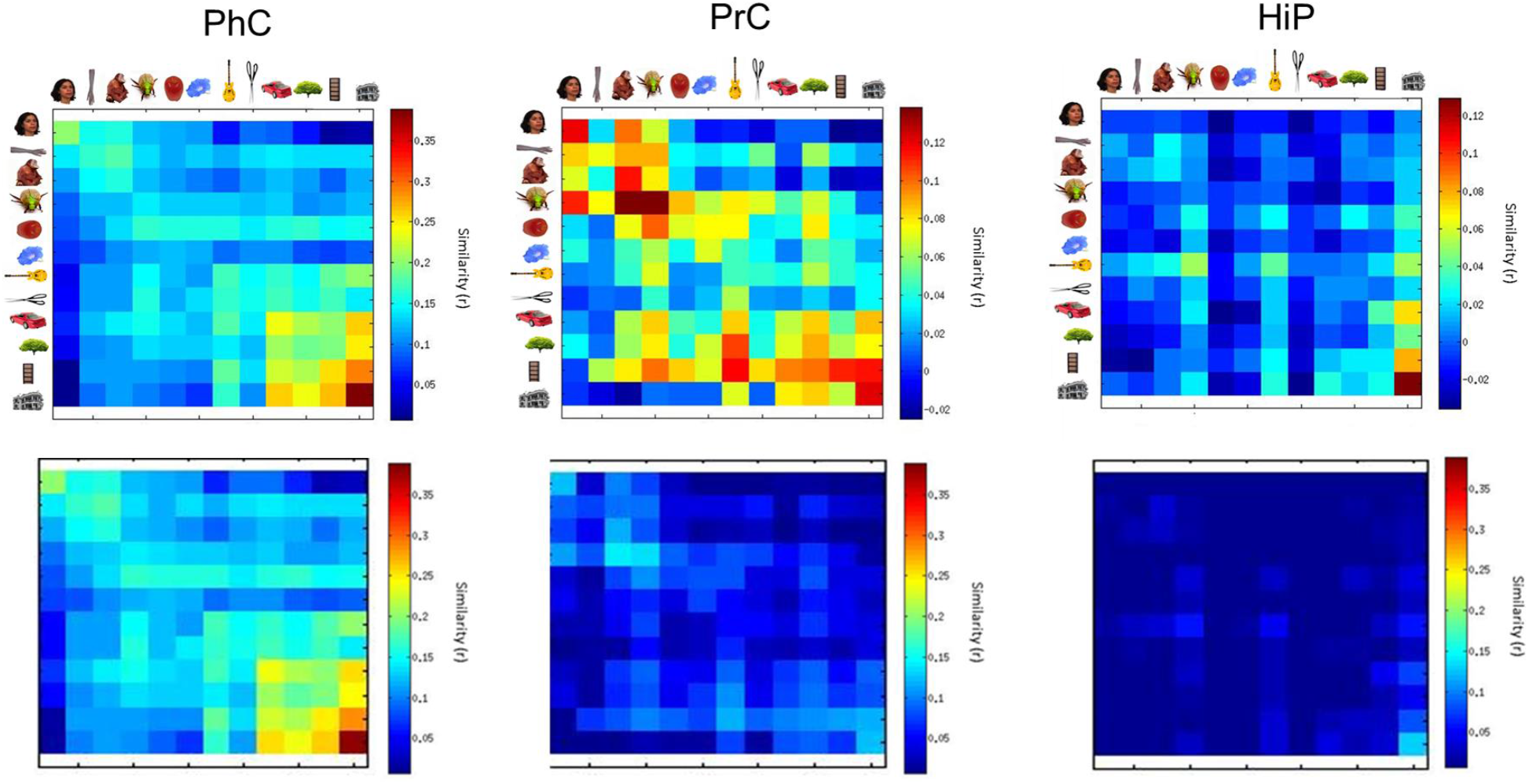
Representational geometry for object evoked responses in the medial temporal lobe. Representational similarity matrices for the three MTL structures. Matrices show Pearson’s correlations between patterns of activity evoked by each object category compared to each other object category. Note that the diagonal shows within-category correlations across runs (each run had different exemplars from the given category). The top row shows each RSM without scaling, the bottom row shows each RSM on the same scale. Note that RSM’s are not symmetrical in the visualization, this is because the upper half of the matrix shows the mean from a subset of across run correlations (i.e., cell 1,2 is condition 1 in the even runs correlated with condition 2 in the odd runs, whereas cell 2,1, is condition 1 in the odd runs correlated with condition 2 in the even runs).

### fMRI Results

#### Category

We first tested a model that probed for the presence of category-specific information by comparing within versus between category similarity across all categories combined, we employed Bonferroni correction for the number of ROIs (3) (Fig. 6). We found that all MTL regions showed sensitivity to category membership (PhC: *t*(12) = 6.41, *p* =. 00006; PrC: *t*(12) =5.01, *p* =. 0006; HiP: *t*(12) = 3.67, *p* =. 009 (Fig. 6). Next we examined sensitivity to information about each category individually, asking for each category whether the within pattern similarity for that category (across runs) was more similar than the between pattern similarity (for that category compared to all other tested categories across runs). To adjust for the larger number of corresponding comparisons, we employed Bonferroni correction in these analyses. In PhC, we found significant effects for buildings (*t*(12) = 5.62, *p* =. 001), furniture (*t*(12) = 3.85, *p* =.02), vehicles (*t*(12) = 4.15, *p* =. 01/, and faces (*t*(12) = 4.23, *p* =. 01). In PrC we found category related effects for monkeys (*t*(12) = 4.28, *p* = 0.01), and a trend towards significance for faces (*t*(12) = 3.17, *p* = 0.08, uncorrected p = 0.007). In the HiP, we only found one category that showed a trend towards significance, namely buildings (*t*(12) = 3.37, *p* =. 06, uncorrected *p* = 0.005).

**Fig 6.**
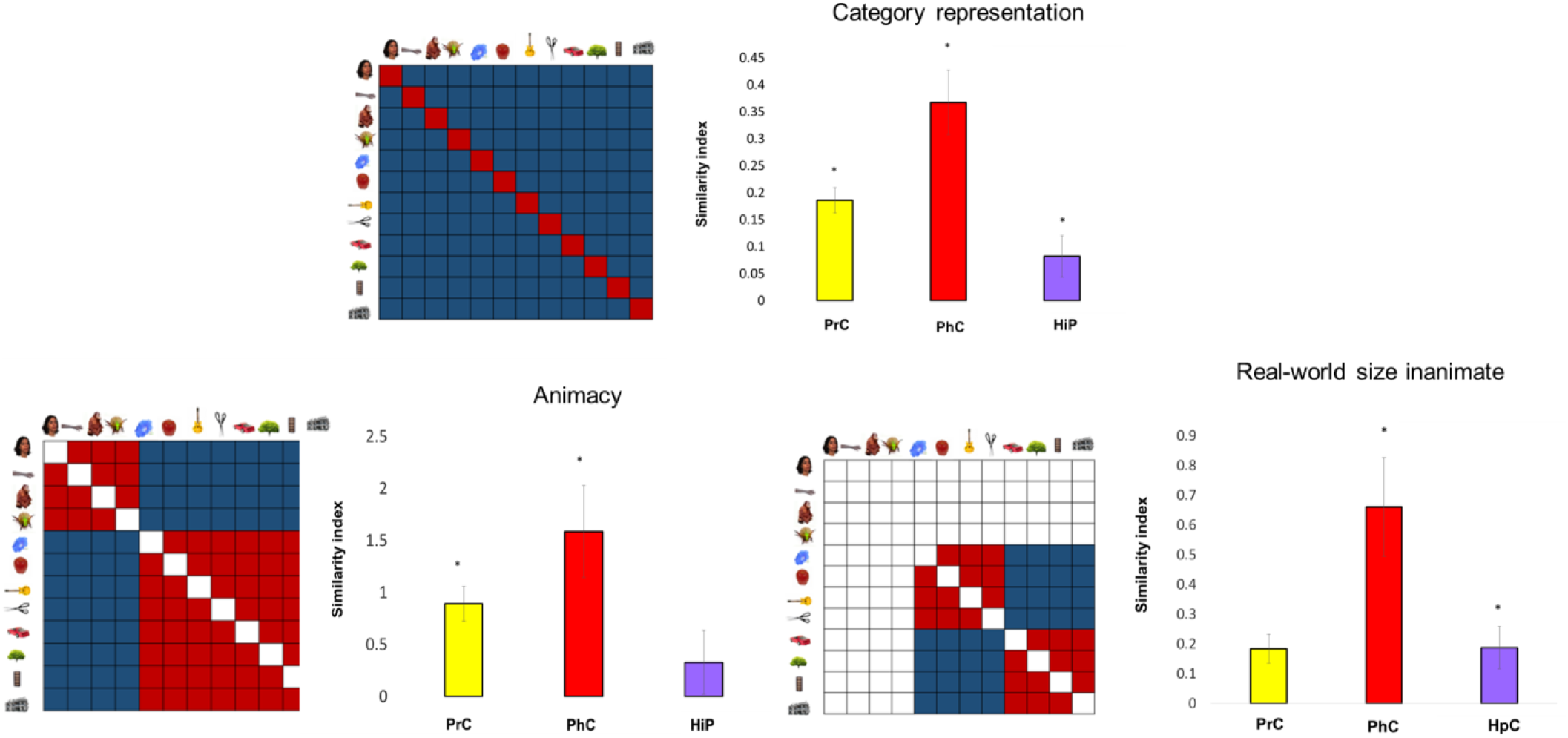
Organization of object representations in the MTL. All bar plots show beta fits between model of organization tested and RSM for each MTL structure. a) model of category representation b) model of animacy organization c) model of real-world size for the inanimate domain.* indicates the model fit was significant with correction for multiple comparisons.

**Fig 7.**
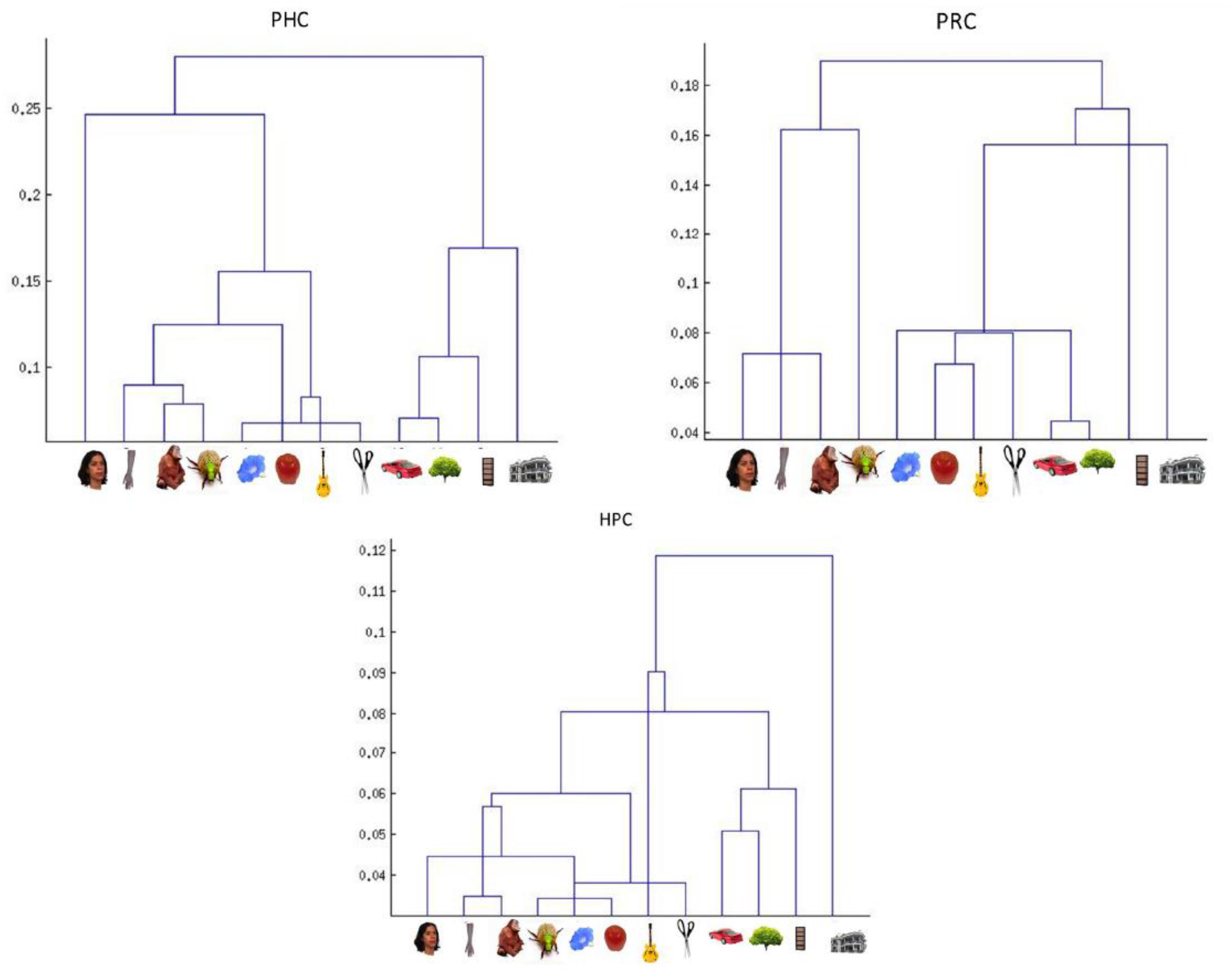
Visualization of representational space. Hierarchical clustering for all object categories in each MTL structure.

#### Animacy

In our next set of analyses we turned to domain-level organization of object representations based on groupings of multiple categories. Specifically, we asked whether MTL regions hold information shared between categories at the domain level of animacy. To address this question, we probed whether representations for objects within a domain (animate or inanimate, respectively) share more similarity with each other than they do with representations from the other domain. In order to remove any impact of category-level effects (as described in the previous section), we removed the diagonal in this model (Fig. 6). We found that the representational structure in both PhC and PrC reflected the animacy divide (PhC: *t*(12) = 3.73, *p* =. 002; PrC: *t*(12)= 3.02, *p* =. 02). By contrast, we found no evidence for organization of object representations by animacy in the HiP (*t*(12) = 2.04, *p* = 0.18) (Fig. 6).

Because there is evidence suggesting that PrC is sensitive to feature overlap, and feature overlap is known to differ across natural kinds versus artifacts (McRae et al., 1997; Devlin et al., 1998; Moss et al., 1998; Tyler et al., 2000; Tyler & Moss, 2001; McRae and Cree, 2002), we also explored whether representations in PrC are organized according to a natural versus artifact divide. Specifically, we compared the categories of flowers, fruits, and trees, with furniture, tools, vehicles, and buildings. The outcome of this analysis, however, provided no evidence in support of this domain organization in PrC (*t*(12) = 1.97, *p* = 0.21).

#### Real-world size

In a further set of analyses, we examined domain-level organization related to the size of inanimate objects. To address this question, we probed whether representations for objects within the domain of small or large inanimate objects, respectively, share more similarity with each other than they do with representations from the other domain. Again, we removed the diagonal in this model in order to remove any impact of category-level effects (Fig. 6). We found evidence for size related organization in both the PhC and HiP (PhC: *t*(12) = 4.14, *p* =. 003; HiP: t(12) = 4.07, p =. 003). By contrast we found no such evidence in PrC (*t*(12) =2.67, *p* = 0.06) (Fig. 6).

#### Visualization of Representational Geometry

In a final step we visualized the representational space for all object categories in each of the ROIs examined using hierarchical clustering (Fig 7.). This data driven approach can be useful in that it can reveal properties that drive the organization of representations without any a priori hypotheses (Kriegeskorte et al., 2008). In PhC, the most dominant dimension of organization is that between large inanimate objects and all other categories. In PrC, the most dominant dimension of organization is animacy. Unlike in PhC, large inanimate objects do not form a separate grouping. Finally, in HiP the most notable distinction is that between buildings and all other object categories.

#### Comparing domain organization within and between MTL structures

Given that we see a different pattern of significant model fits across MTL ROI’s, to further probe these differences we compared model fits for different domains to each other within each ROI. Specifically, we tested whether organization by real-world size or animacy was a better fit within each ROI by computing a within-subjects t-test on the beta-values for the model fits. We found no significant differences between model fits within any ROI (PhC: *t*(12) = 0.46, *p* = 0.65; PrC: *t*(12) = 0.56, *p* = 0.58; HiP: *t*(12) = 0.71, *p* = 0.48). We also asked whether organization by category or domain differed significantly across region when tested against each other. Specifically, we ran a repeated measures ANOVA with ROI (PhC, PrC, HiP) and model (category, animacy, real-world size for inanimate) as factors. We found a main effect of ROI *F*(12) = 4.587, *p* = 0.04, but no interaction (roi × model) *F*(12) = 0.026, *p* = 0.97.

#### Additional analyses

Our main goal in the current study was to explore representational space in MTL structures. However, we completed two further analyses to better understand the selectivity of our MTL findings. First, we tested all three models (category membership, animacy, and real-world size) in a control region where we would not expect to see organization for visual stimuli, namely primary auditory cortex bilaterally. Second, we tested all three models in two visual cortex ROIs, a bilateral primary visual cortex ROI, and lateral occipital cortex (LOC), in order to compare and contrast these regions with the MTL regions. The primary auditory and visual cortex ROI’s were taken from the MarsBar toolbox (Mathews, 2002), the LOC ROI was taken from Xu et al., 2006. All ROI’s were transformed from MNI space to native space for each subject, and subsequent analyses was identical to that previously described. We found no evidence of category or domain organization in primary auditory cortex (category: *t*(12) = −0.70, *p* = 0.50; animacy: *t*(12) = −0.23,*p* = 0.82; real-world size: *t*(12) = −0.60,*p* = 0.55, *uncorrected*). In primary visual cortex, all three models were significant (category: *t*(12) = 8.27, *p* = 0.00001; animacy: *t*(12) = 6.05, *p* = 0.0002; real-world size: *t*(12), *p* = 0.005). In LOC, we found category organization (*t*(12) = 8.5, *p* = 0.00001) as well as animacy (*t*(12) = 3.65, *p* = 0.01), but not real-world size (*t*(12) = 1.76, *p* =0.31).

## DISCUSSION

In the current study we examined the organization of object representations in MTL structures, aiming to determine whether dimensions of organization prominent in upstream VVS are present in the MTL when participants perform a recognition-memory task. Specifically, we asked (i) whether there is category specificity in object representations in MTL structures (i.e., PrC, PhC, HiP), (ii) whether there is domain specificity along an animate-inanimate divide, and (iii) whether there is specificity in representations for inanimate objects related to real-world size. We found that similar to VVS representational organization, MTL structures do indeed display sensitivity to category membership, animacy, and real-world size for inanimate objects. While model fits related to these dimensions differed across structures when probed individually, hinting at interesting links to functional differentiation previously discussed in the literature, similarities in organization across MTL structures were more prominent overall. Our findings replicate and extend previous findings pertaining to category specificity in other task contexts. Critically, they also expand the extant literature by providing first insight into domain-level organization of object representations in the MTL during recognition memory.

### PrC

PrC is the MTL structure that has most extensively been linked to object processing in prior research. While this has been best characterized with respect to its role in recognition memory for objects, recent work suggests that object representations in PrC also play a critical role in perceptual and semantic tasks (Barense et al., 2010; Bussey et al., 2002; Kivisaari et al., 2012, Clarke & Tyler, 2014; Bruffaerts et al., 2013; Martin et al., 2018; Lsee Graham et al., 2010, for review). However, the organization of object representations that support judgements in these tasks has received only limited investigation so far. In terms of category-level organization, it has been reported that PrC shows specificity for the category of faces in recognition memory and perceptual tasks (Diana et al., 2007; Martin et al., 2013, 2016; O’Neil et al. 2013, 2014). The present result extend this prior research by showing that PrC also shows specificity for another animate category, namely monkeys, in combination with a trend towards specificity for faces. Beyond this category-level organization, we see a broader organization in PrC by the domain of animacy.

To our knowledge, domain-level organization has only been explored previously in tasks that require object naming at the basic (rather than exemplar) level. Specifically, it has been reported that PrC shows higher levels of activity when participants have to name objects that are animate as compared to objects that are inanimate (Moss et al., 2005) and there is also evidence that damage to the PrC differentially affects naming for animate objects (Wright et al., 2015). This domain-specific pattern of findings has been attributed to the fact that animate objects are distinct from inanimate objects at the level of feature statistics. Specifically, one important dimension that differs across animate and inanimate objects is the amount of feature overlap and feature distinctiveness amongst members of those domains. It has been argued that overall animate objects have more feature overlap and less distinctive features than inanimate objects (McRae et al., 1997; Devlin et al., 1998; Moss et al., 1998; Tyler et al., 2000; Tyler & Moss, 2001; McRae and Cree, 2002). In these studies feature overlap is typically defined based on listed features that can be classified as perceptual or semantic (Martin et al., 2018), and the level of representations tapped into by naming are at the basic level (i.e., distinguishing a horse from a zebra rather than two different horses from each other). Indeed, an fMRI study that employed RSA to examine object representations in PrC during naming revealed that PrC uniquely holds information at the individual object level (Clarke & Tyler, 2014; see also Bruffaerts et al., 2013, Martin 2018 for related findings in PrC based on written words). In the context of the continuous recognition memory task used in the current study, participants were required to make discriminations similar, if not more fine-grained, to those required for naming an individual exemplar. Namely, the task required recognition of prior occurrence of specific exemplars, such as whether a particular building had been presented previously. Thus, although our study did not aim to test specific hypotheses about the impact of feature overlap on representational similarities, one possibility is that the animacy-related organization we report reflects differences on this dimension between the animate and inanimate objects we employed. Given that natural but inanimate object categories (such as fruits and vegetables) are also known to have higher feature overlap than artifacts (such as tools and buildings) (McRae et al., 1997; Devlin et al., 1998; Moss et al., 1998; Tyler et al., 2000; Tyler & Moss, 2001; McRae and Cree, 2002), we additionally explored whether PrC might show domain-level organization related to whether an object is natural or an artifact. This analysis, however, did not provide evidence for such a distinction. However, given that this could be due to a lower degree of feature overlap in our natural stimuli subset than our animate subset, further research with explicit modeling of response patterns based on quantitative estimates of feature overlap is required in order to determine how feature overlap contributes to the domain level organization we report here.

The sensitivity of PrC to the animate-inanimate distinction may also relate to the long range connectivity it maintains with other cortical and subcortical regions. The idea that large scale connectivity may drive differential sensitivity between stimuli of different domains, such as animate or large inanimate objects, has been fruitful towards understanding VVS organization in more posterior regions. Using a data-driven approach with resting-state fMRI connectivity data, Konkle & Caramazza (2016) identified three distinct resting state networks that ‘route through’ the large domain-preferring tripartite regions of VVS. Specifically, animate-object preferring regions were more strongly coupled with the anterior temporal lobe, small inanimate-object preferring regions were more strongly coupled with aspects of parietal cortex, and large inanimate object preferring regions were more correlated with the posterior medial temporal lobe, as well as early visual cortex regions differentially involved in processing stimuli in the peripheral visual fields. Current evidence linking long range connectivity in PrC to processing information from particular object domains or categories is very limited at present. However, in a recent study diffusion tensor imaging study microstructure of the inferior longitudinal fasciculus, which connects the occipital and ventro-anterior temporal lobe, including PrC, specifically correlated with accuracy on a perceptual discrimination task involving faces but not scenes, as well category BOLD response to faces in this task (Hodgetts et al., 2015). In addition, several studies have examined the resting state connectivity profiles that characterize different MTL structures (Kahn et al., 2008; Libby et al., 2012). At the whole brain level, PrC shows distinct connectivity with other structures within the anterior temporal lobes, amygdala, and lateral orbitofrontal cortex. These connectivity findings have led to the suggestion the PrC is part of a cortical network, referred to as the anterior-temporal network that plays a unique functional role in memory and cognition (Ranganath & Ritchey, 2012). It has been argued that, relative to a posterior-medial system of which PhC is a central component, this anterior system is preferentially involved in object recognition as well as processing the social and emotional aspects of objects and animate entities, semantic knowledge, and reward learning. Although the model does not explicitly consider differences between specific object categories or domains, to the extent that the information processed in the anterior system pertains to ecologically relevant information, this kind of processing may be more relevant to animate objects.

### PhC

The role of the PhC in object processing during naming and recognition memory tasks has been less explored than that of PrC, including evaluating any function of feature overlap. In the memory literature, PhC has been primarily implicated in scene recognition and in context representation in tasks of associative memory (Ranganath & Ritchey, 2012). However, recently, it has been shown that PhC also plays a role in recognition memory for objects, specifically those that have navigational relevance, such as buildings or trees (Martin et al., 2013; Martin et al., 2018; see also Janzen & van Turennout, 2004). In the current study, we also found category specificity for buildings and trees, in addition to other large inanimate objects, including furniture and vehicles. At the domain level, we observed organization by animacy and real-world size for inanimate objects. This is notable because the PPA (or parahippocampal place area), which includes the posterior portion of PhC, has also been shown to have higher levels of activity for inanimate objects, even when contrasted with shape-matched animate objects (Proklova, Kaiser, and Peelen, 2016). Moreover, a number of studies have demonstrated that the PPA is more active for large than for small objects (Konkle et al., 2013; Aguirre et al, 1998; Julian et al., 2016), and most similar to our findings, that patterns of activity in the PPA distinguish between large and small objects (Julian et al., 2016). This sensitivity to real-world size in the PPA, as well as that in the PhC more broadly that we describe here, appears to be more reliable than what is observed in PrC, where it reflected only a trend in the current study. This pattern could suggest that there may be a gradient in terms of coding for real-world size along the anterior-posterior axis of the parahippocampal gyrus.

As in our discussion pertaining to PrC, it is informative to consider the long-range connectivity of PhC in relation to the category and domain level organization reported here. Resting state connectivity studies at the whole brain level have shown that PhC is differentially connected to the retrosplenial cortex (RSC), posterior cingulate, precuneus, parietal cortex, and ventromedial prefrontal cortex, the thalamus. In addition, PhC is also more strongly connected to posterior medial occipital cortex as well as early visual areas (Libby et al., 2012). In light of these resting-state connectivity findings, it has been suggested that PhC is a component of the posterior medial network, with a functional role in memory and cognition that differs from that of the anterior-temporal network that includes PrC. These findings generally align with the findings reported by Konkle et al. (2016) that cortex in the medial VVS that prefers large inanimate object is highly connected to early visual areas tuned to the peripheral visual fields as well as MTL (although not clearly specified whether it is the posterior portion of the parahippocampal gyrus, it is distinct from the anterior temporal area more highly connected to lateral animate VVS cortex). It has been argued that this network is important for representing context in episodic memory and episodic simulation, as well as in spatial navigation (Ranganath & Ritchey, 2012). One possibility is that the sensitivity of PhC to the animacy divide we report here is linked to differential processing of large inanimate objects that are important for navigation, or are more likely to serve as episodic context. Compared to animate objects, large inanimate objects often evoke a stronger sense of surrounding space (Mullally & Maguire, 2011), and when stable, can also serve as landmarks (Martin et al. under review; Janzen & van Turennout, 2004; Troiani et al., 2013). From this perspective, animacy plays a role in the organization of object representations in PhC because large inanimate objects share dimensions important for the general functions of a posterior-medial cortical system. We note, however, that any such preferential role does not appear to be absolute as PhC also appears to represent faces as a distinct category as observed in the current study and in other prior research (Huffman & Stark, 2014; Diana et al., 2008; Liang et al., 2012). At a more general level, such findings suggest that the organization of object representations in the MTL also resembles that in the VVS, by virtue of pointing to distributed representations crossing multiple structures rather than sharply defined functional modules (Haxby et al., 2001).

### HiP

Interestingly, we found that the HiP shows no clear-cut categorical representations of objects, although we observed a trend for the representation of buildings, or organization by animacy. Similar to PhC, the HiP was sensitive to the distinction between large and small inanimate objects. The lack of clear cut category-specific representation in our findings is in line with previous suggestions that the HiP is agnostic to the nature or content of its representations at the item level. The agnosticity of the HiP has been attributed to its unique role in pattern separation of episodes (Huffman & Stark, 2014). According to this reasoning, the result of hippocampal pattern separation is that representations in the HiP are more dissimilar to each other than those in PrC and PhC, leading to the loss of specificity in organization by category that is present in these input structures. However, the evidence for domain-level organization related to size we report here suggests that the HiP may not be entirely insensitive to content.

There is substantial evidence for a role of the HiP in scene perception and construction (Hodgetts et al., 2016; Lee et al., 2005; Barense et al., 2015; Zeidman et al., 2015; for review see Murray et al., 2017). For example, it has been demonstrated that the HiP is more active during perceptual oddity tasks for scenes than for other types of stimuli (Lee et al., 2008). Hodgetts et al. (2016) found clusters of activity in the HiP that are higher for scenes than for other stimulus categories (faces, objects) while participants performed a 1-back task, and these clusters appeared as reliably as clusters in the traditional scene-processing network (including PhC, retrosplenial cortex, and transverse occipital sulcus). Based on these results, the authors suggested that the HiP should be considered as a component of the core scene processing network. Implied with this argument is the notion that the HiP is not entirely agnostic to stimulus content. More recent work by this group of researchers has provided some evidence to explain why some studies find evidence for differential involvements in scene processing and other do not (Hodgetts et al., 2017). In that fMRI study, conducted with ultra-high resolution, sensitivity to scene stimuli could be more precisely localized to a specific subfield of the HiP, namely the subiculum, with other subfields staying agnostic. It is possible that the sensitivity to real-world size of objects reported here, together with the hint for category specific representations for buildings in HiP, are a result of similarities between large objects and scenes that are of particular relevance to processing in the subiculum.

## Conclusions

Together, our findings show that stimulus dimensions that influence the organization of object representations in the VVS also shape this organization in the MTL. Moreover, they reveal many similarities in organization across PrC, PhC, and the HiP, with only some hints of differences. A promising direction for future research will be to test for these differences in a more targeted manner. In addition, it will be important to examine how patterns of large-scale connectivity can account for the organizational principles in the MTL described, and to determine how they relate to specific functional and perceptual properties of objects that differ across domains and categories.

---

| category | d' | RT cr's | RT hits | Sig differences overall RT | Sig differences dprime |
| --- | --- | --- | --- | --- | --- |
| face | 1.58, sd=0.35 | 1.037, sd=0.041 | .975 sd=0.057 |  |  |
| body | 1.60, sd=0.61 | 1.059, sd=0.042 | .979, sd=0.045 | fruit, tool |  |
| monkey | 1.69, sd=0.66 | 1.056, sd=0.047 | .980, sd=0.034 | fruit, tool |  |
| insect | 2.18, sd=0.97 | 1.041 sd=0.038 | .972, sd=0.046 |  |  |
| flower | 1.93, sd=0.86 | 1.017, sd=0.027 | .991, sd=0.070 |  |  |
| fruit | 2.33, sd=0.95 | 1.023, sd=0.045 | .951, sd=0.038 | tree, building | tree |
| musical instrument | 1.88, sd=0.59 | 1.012, sd=0.034 | .968, sd=0.046 |  |  |
| tool | 1.90, sd=0.58 | 1.014, sd=0.048 | .965, sd=0.047 | building |  |
| tree | 1.62, sd=0.71 | 1.035, sd=0.042 | .994, sd=0.042 |  |  |
| vehicle | 2.51, sd=0.86 | 1.007, sd=0.047 | .987, sd=0.063 |  | body, face, fruit, furniture, tree, monkey |
| furniture | 1.71, sd=0.64 | 1.029, sd=0.042 | .974, sd=0.040 |  |  |
| building | 1.99, sd=0.72 | 1.036, sd=0.048 | .995, sd=0.049 |  |  |

## References

Aguirre, G. K., Zarahn, E., & D’esposito, M. (1998). An area within human ventral cortex sensitive to “building” stimuli: evidence and implications. Neuron, 21(2), 373–383.

Barense, M. D., Henson, R. N., Lee, A. C., & Graham, K. S. (2010). Medial temporal lobe activity during complex discrimination of faces, objects, and scenes: Effects of viewpoint. Hippocampus, 20(3), 389–401.

Bussey, T. J., & Saksida, L. M. (2005). Object memory and perception in the medial temporal lobe: an alternative approach. Current opinion in neurobiology, 15(6), 730–737.

Bussey, T. J., & Saksida, L. M. (2007). Memory, perception, and the ventral visual-perirhinal-hippocampal stream: thinking outside of the boxes. Hippocampus, 17(9), 898–908.

Bruffaerts, R., Dupont, P., Peeters, R., De Deyne, S., Storms, G., & Vandenberghe, R. (2013). Similarity of fMRI activity patterns in left perirhinal cortex reflects semantic similarity between words. Journal of Neuroscience, 33(47), 18597–18607.

Bussey, T. J., Saksida, L. M., & Murray, E. A. (2002). Perirhinal cortex resolves feature ambiguity in complex visual discriminations. European Journal of Neuroscience, 15(2), 365–374.

Capitani, E., Laiacona, M., Mahon, B. Z., & Caramazza, A. (2003). What are the facts of semantic category-specific deficits: A critical review of the clinical evidence. Cognitive Neuropsychology, 20, 213–261.

Caramazza, A., & Mahon, B. Z. (2003). The organization of conceptual knowledge: the evidence from category-specific semantic deficits. Trends in cognitive sciences, 7(8), 354–361.

Cohen, L., & Dehaene, S. (2004). Specialization within the ventral stream: the case for the visual word form area. Neuroimage, 22(1), 466–476.

Clarke, A., & Tyler, L. K. (2014). Object-specific semantic coding in human perirhinal cortex. The Journal of Neuroscience, 34(14), 4766–4775.

Cusack R, Vicente-Grabovetsky A, Mitchell DJ, Wild CJ, Auer T, Linke AC, Peelle JE (2015) Automatic analysis (aa): Efficient neuroimaging workflows and parallel processing using Matlab and XML. Frontiers in Neuroinformatics 8:90.http://dx.doi.org/10.3389/fninf.2014.00090.

Davachi L. (2006). Item, context and relational episodic encoding in humans. Current opinion in neurobiology, 16(6), 693–700.

Damasio, A. R., Damasio, H., & Van Hoesen, G. W. (1982). Prosopagnosia Anatomic basis and behavioral mechanisms. Neurology, 32(4), 331–331.

De Beeck, H. P. O., Haushofer, J., & Kanwisher, N. G. (2008). Interpreting fMRI data: maps, modules and dimensions. Nature reviews. Neuroscience, 9(2), 123.

De Renzi, E., Perani, D., Carlesimo, G. A., Silveri, M. C., & Fazio, F. (1994). Prosopagnosia can be associated with damage confined to the right hemisphere—an MRI and PET study and a review of the literature. Neuropsychologia, 32(8), 893–902.

Devlin, J. T., Gonnerman, L. M., Andersen, E. S., & Seidenberg, M. S. (1998). Category-specific semantic deficits in focal and widespread brain damage: A computational account. Journal of cognitive Neuroscience, 10(1), 77–94.

Downing, P. E., Jiang, Y., Shuman, M., & Kanwisher, N. (2001). A cortical area selective for visual processing of the human body. Science, 293(5539), 2470–2473.

Epstein, R., & Kanwisher, N. (1998). A cortical representation of the local visual environment. Nature, 392(6676), 598–601.

Epstein, R., Harris, A., Stanley, D., & Kanwisher, N. (1999). The parahippocampal place area: recognition, navigation, or encoding?. Neuron, 23(1), 115–125.

Eichenbaum, H., Yonelinas, A. P., & Ranganath, C. (2007). The medial temporal lobe and recognition memory. Annu. Rev. Neurosci., 30, 123–152.

Epstein, R. A., & Vass, L. K. (2014). Neural systems for landmark-based wayfinding in humans. Phil. Trans. R. Soc. B, 369(1635), 20120533.

Frankó, E., Insausti, A. M., Artacho-Pérula, E., Insausti, R., & Chavoix, C. (2014). Identification of the human medial temporal lobe regions on magnetic resonance images. Human brain mapping, 35(1), 248–256.

Graham, K. S., Barense, M. D., & Lee, A. C. (2010). Going beyond LTM in the MTL: a synthesis of neuropsychological and neuroimaging findings on the role of the medial temporal lobe in memory and perception. Neuropsychologia, 48(4), 831–853.

Grill-Spector, K., & Weiner, K. S. (2014). The functional architecture of the ventral temporal cortex and its role in categorization. Nature Reviews Neuroscience, 15(8), 536–548.

Guderian, S., Dzieciol, A. M., Gadian, D. G., Jentschke, S., Doeller, C. F., Burgess, N., … & Vargha-Khadem, F. (2015). Hippocampal volume reduction in humans predicts impaired allocentric spatial memory in virtual-reality navigation. Journal of Neuroscience, 35(42), 14123–14131.

Hart, J., Berndt, R. S., & Caramazza, A. (1985). Category-specific naming deficit following cerebral infarction. Nature, 316(6027), 439–440.

Hassabis, D., Kumaran, D., Vann, S. D., & Maguire, E. A. (2007). Patients with hippocampal amnesia cannot imagine new experiences. Proceedings of the National Academy of Sciences, 104(5), 1726–1731.

Haxby, J. V., Gobbini, M. I., Furey, M. L., Ishai, A., Schouten, J. L., & Pietrini, P. (2001). Distributed and overlapping representations of faces and objects in ventral temporal cortex. Science, 293(5539), 2425–2430.

Hillis, A. E., & Caramazza, A. (1991). Category-specific naming and comprehension impairment: A double dissociation. Brain, 114(5), 2081–2094.

Hodgetts, C. J., Shine, J. P., Lawrence, A. D., Downing, P. E., & Graham, K. S. (2016). Evidencing a place for the hippocampus within the core scene processing network. Human brain mapping, 37(11), 3779–3794.

Huffman, D. J., & Stark, C. E. (2014). Multivariate pattern analysis of the human medial temporal lobe revealed representationally categorical cortex and representationally agnostic hippocampus. Hippocampus, 24(11), 1394–1403.

Humphreys, G. W., & Riddoch, M. J. (2003). A case series analysis of “category-specific” deficits of living things: The HIT account. Cognitive Neuropsychology, 20(3-6), 263–306.

Janzen, G., & Van Turennout, M. (2004). Selective neural representation of objects relevant for navigation. Nature neuroscience, 7(6), 673.

Johnson-Frey A. H. (2004). The neural basis of complex tool use in humans. Trends in Cognitive Sciences, 8(2), 71–78.

Julian, J. B., Ryan, J., & Epstein, R. A. (2016). Coding of Object Size and Object Category in Human Visual Cortex. Cerebral Cortex, bhw150.

Kanwisher, N., McDermott, J., & Chun, M. M. (1997). The fusiform face area: a module in human extrastriate cortex specialized for face perception. The Journal of neuroscience, 17(11), 4302–4311.

Kanwisher, N., & Dilks, D. D. (2013). The functional organization of the ventral visual pathway in humans. The new visual neurosciences, 733–748.

Konkle, T., & Oliva, A. (2012). A real-world size organization of object responses in occipitotemporal cortex. Neuron, 74(6), 1114–1124.

Konkle, T., & Caramazza, A. (2013). Tripartite organization of the ventral stream by animacy and object size. The Journal of Neuroscience, 33(25), 10235–10242.

Konkle, T., & Caramazza, A. (2016). The large-scale organization of object-responsive cortex is reflected in resting-state network architecture. Cerebral Cortex, 1–13.

Kivisaari, S. L., Tyler, L. K., Monsch, A. U., & Taylor, K. I. (2012). Medial perirhinal cortex disambiguates confusable objects. Brain, 135(12), 3757–3769.

Kriegeskorte, N., Mur, M., & Bandettini, P. A. (2008). Representational similarity analysis-connecting the branches of systems neuroscience. Frontiers in systems neuroscience, 2, 4.

Kriegeskorte, N., Mur, M., Ruff, D. A., Kiani, R., Bodurka, J., Esteky, H., … Bandettini, P. A. (2008). Matching categorical object representations in inferior temporal cortex of man and monkey. Neuron, 60(6), 1126–1141.

Lee, A. C., Scahill, V. L., & Graham, K. S. (2007). Activating the medial temporal lobe during oddity judgment for faces and scenes. Cerebral Cortex, 18(3), 683–696.

Lee, A. C., Buckley, M. J., Pegman, S. J., Spiers, H., Scahill, V. L., Gaffan, D., … & Graham, K. S. (2005). Specialization in the medial temporal lobe for processing of objects and scenes. Hippocampus, 15(6), 782–797.

Liang, J. C., Wagner, A. D., & Preston, A. R. (2013). Content representation in the human medial temporal lobe. Cerebral Cortex, 23(1), 80–96.

Libby, L. A., Ekstrom, A. D., Ragland, J. D., & Ranganath, C. (2012). Differential connectivity of perirhinal and parahippocampal cortices within human hippocampal subregions revealed by high-resolution functional imaging. Journal of Neuroscience, 32(19), 6550–6560.

Linke, A. C., Vicente-Grabovetsky, A., & Cusack, R. (2011). Stimulus-specific suppression preserves information in auditory short-term memory. Proceedings of the National Academy of Sciences, 108(31), 12961–12966.

Litman, L., Awipi, T., & Davachi, L. (2009). Category-specificity in the human medial temporal lobe cortex. Hippocampus, 19(3), 308–319.

Martin A. (2007). The representation of object concepts in the brain. Annu. Rev. Psychol., 58, 25–45.

Martin, C. B., McLean, D. A., O’Neil, E. B., & Köhler, S. (2013). Distinct familiarity-based response patterns for faces and buildings in perirhinal and parahippocampal cortex. The Journal of Neuroscience, 33(26), 10915–10923.

Martin, C. B., Cowell, R. A., Gribble, P. L., Wright, J., & Köhler, S. (2015). Distributed category-specific recognition-memory signals in human perirhinal cortex. Hippocampus.

Martin, C. B., Sullivan, J. A., Wright, J., & Köhler, S. (2018). How landmark suitability shapes recognition memory signals for objects in the medial temporal lobes. NeuroImage, 166, 425–436.

Martin, C. B., Douglas, D., Newsome, R. N., Man, L. L., & Barense, M. D. (2018). Integrative and distinctive coding of visual and conceptual object features in the ventral visual stream. Elife, 7, e31873.

Matthew Brett, Jean-Luc Anton, Romain Valabregue, Jean-Baptiste Poline. Region of interest analysis using an SPM toolbox [abstract] Presented at the 8th International Conference on Functional Mapping of the Human Brain, June 2-6, 2002, Sendai, Japan. Available on CD-ROM in NeuroImage, Vol 16, No 2.

McRae, K., De Sa, V. R., & Seidenberg, M. S. (1997). On the nature and scope of featural representations of word meaning. Journal of Experimental Psychology: General, 126(2), 99.

McRae, K., & Cree, G. S. (2002). Factors underlying category-specific semantic deficits. Category-specificity in brain and mind, 211–249.

Moss, H. E., Tyler, L. K., Durrant-Peatfield, M., & Bunn, E. M. (1998). ‘Two eyes of a see-through’: Impaired and intact semantic knowledge in a case of selective deficit for living things. Neurocase, 4(4-5), 291–310.

Moss, H. E., Rodd, J. M., Stamatakis, E. A., Bright, P., & Tyler, L. K. (2004). Anteromedial temporal cortex supports fine-grained differentiation among objects. Cerebral cortex, 15(5), 616–627.

Mullally, S. L., Intraub, H., & Maguire, E. A. (2012). Attenuated boundary extension produces a paradoxical memory advantage in amnesic patients. Current Biology, 22(4), 261–268.

Murray, E. A., Wise, S. P., & Graham, K. S. (2017). Representational specializations of the hippocampus in phylogenetic perspective. Neuroscience Letters.

O’Neil, E. B., Barkley, V. A., & Köhler, S. (2013). Representational demands modulate involvement of perirhinal cortex in face processing. Hippocampus, 23(7), 592–605.

O’Neil, E. B., Hutchison, R. M., McLean, D. A., & Köhler, S. (2014). Resting-state fMRI reveals functional connectivity between face-selective perirhinal cortex and the fusiform face area related to face inversion. Neuroimage, 92, 349–355.

Proklova, D., Kaiser, D., & Peelen, M. V. (2016). Disentangling representations of object shape and object category in human visual cortex: The animate-inanimate distinction. Journal of cognitive neuroscience.

Pruessner, J. C., Li, L. M., Serles, W., Pruessner, M., Collins, D. L., Kabani, N., … & Evans, A. C. (2000). Volumetry of hippocampus and amygdala with high-resolution MRI and threedimensional analysis software: minimizing the discrepancies between laboratories. Cerebral cortex, 10(4), 433–442.

Pruessner, J. C., Köhler, S., Crane, J., Pruessner, M., Lord, C., Byrne, A., … & Evans, A. C. (2002). Volumetry of temporopolar, perirhinal, entorhinal and parahippocampal cortex from high-resolution MR images: considering the variability of the collateral sulcus. Cerebral Cortex, 12(12), 1342–1353.

Ralph, M. A. L., Lowe, C., & Rogers, T. T. (2007). Neural basis of category-specific semantic deficits for living things: evidence from semantic dementia, HSVE and a neural network model. Brain, 130(4), 1127–1137.

Sacchett, C., & Humphreys, G. W. (1992). Calling a squirrel a squirrel but a canoe a wigwam: A category-specific deficit for artefactual objects and body parts. Cognitive Neuropsychology, 9(1), 73–86.

Scaplen, K. M., Gulati, A. A., Heimer-McGinn, V. L., & Burwell, R. D. (2014). Objects and landmarks: hippocampal place cells respond differently to manipulations of visual cues depending on size, perspective, and experience. Hippocampus, 24(11), 1287–1299.

Sha, L., Haxby, J. V., Abdi, H., Guntupalli, J. S., Oosterhof, N. N., Halchenko, Y. O., & Connolly, A. C. (2015). The animacy continuum in the human ventral vision pathway. Journal of cognitive neuroscience.

Schacter, D. L., Addis, D. R., & Buckner, R. L. (2007). Remembering the past to imagine the future: the prospective brain. Nature reviews. Neuroscience, 8(9), 657.

Troiani, V., Stigliani, A., Smith, M. E., & Epstein, R. A. (2012). Multiple object properties drive scene-selective regions. Cerebral Cortex, 24(4), 883–897.

Tyler, L. K., Moss, H. E., Durrant-Peatfield, M. R., & Levy, J. P. (2000). Conceptual structure and the structure of concepts: A distributed account of category-specific deficits. Brain and language, 75(2), 195–231.

Tyler, L. K., & Moss, H. E. (2001). Towards a distributed account of conceptual knowledge. Trends in cognitive sciences, 5(6), 244–252.

van der Ham, I. J., Martens, M. A., Claessen, M. H., & van den Berg, E. (2017). Landmark Agnosia: Evaluating the Definition of Landmark-based Navigation Impairment. Archives of Clinical Neuropsychology, 32(4), 472–482.

Walther, A., Nili, H., Ejaz, N., Alink, A., Kriegeskorte, N., & Diedrichsen, J. (2016). Reliability of dissimilarity measures for multi-voxel pattern analysis. Neuroimage, 137, 188–200.

Warrington, E. K., & McCarthy, R. A. (1988). The fractionation of retrograde amnesia. Brain and cognition, 7(2), 184–200.

Warrington, E. K., & Shallice, T. (1984). Category specific semantic impairments. Brain, 107(3), 829–853.

Wright, P., Randall, B., Clarke, A., & Tyler, L. K. (2015). The perirhinal cortex and conceptual processing: effects of feature-based statistics following damage to the anterior temporal lobes. Neuropsychologia, 76, 192–207.

Xu, Y., & Chun, M. M. (2006). Dissociable neural mechanisms supporting visual short-term memory for objects. Nature, 440(7080), 91.

